# High-resolution kinetic gene expression analysis of T helper cell differentiation reveals a STAT-dependent, unique transcriptional program in Th1/2 hybrid cells

**DOI:** 10.1101/2022.05.13.491791

**Authors:** Philipp Burt, Michael Peine, Caroline Peine, Zuzanna Borek, Sebastian Serve, Michael Floßdorf, Ahmed N. Hegazy, Thomas Höfer, Max Löhning, Kevin Thurley

## Abstract

Selective differentiation of CD4+ T helper (Th) cells into specialized subsets such as Th1 and Th2 cells is a key element of the adaptive immune system driving appropriate immune responses. Besides those canonical Th cell lineages, hybrid phenotypes such as Th1/2 cells arise in vivo, and their generation could be reproduced in vitro. While master-regulator transcription factors like T-bet for Th1 and GATA-3 for Th2 cells drive and maintain differentiation into the canonical lineages, the transcriptional architecture of hybrid phenotypes is less well understood. In particular, it has remained unclear whether a hybrid phenotype implies a mixture of the effects of several canonical lineages for each gene, or rather a bimodal behavior across genes. Th cell differentiation is a dynamic process in which the regulatory factors are modulated over time, but longitudinal studies of Th cell differentiation are sparse. Here, we present a dynamic transcriptome analysis following Th cell differentiation into Th1, Th2 and Th1/2 hybrid cells. We identified an early bifurcation point in gene expression programs, and we found that only a minority of ∼20% of Th cell-specific genes showed mixed effects from both Th1 and Th2 cells on Th1/2 hybrid cells. While most genes followed either Th1 or Th2 cell gene expression, another fraction of ∼20% of genes followed a Th1 and Th2 cell-independent transcriptional program under control of the transcription factors STAT1 and STAT4. Overall, our results emphasize the key role of high-resolution longitudinal data for the characterization of cellular phenotypes.

## Introduction

The differentiation of CD4+ T helper (Th) cells into effector cell lineages associated with specific immunological functions is a critical event at the onset of an immune response. Individual Th cell lineages such as Th1 and Th2 cells can be discriminated by expression of the master-regulator transcription factors T-bet and GATA-3, and by production of signature cytokines such as IFN-γ and IL-4, respectively (1,2). The differentiation process from naïve Th cells into the various effector cell lineages spans multiple days, and the underlying transcriptional network governing the decision processes changes dynamically throughout differentiation (3,4). The gene-regulatory networks for Th cell subset-specific differentiation are quite complex and can be modulated by cell-cell interactions (5). Th cell phenotypes are not limited to the canonical Thx phenotypes (Th1, Th2, Th17,amongst others), but also include stable hybrid forms such as Th1/2 cells, which co-express T-bet and GATA-3 as well as IFN-γ and IL-4 (6–8).

In previous studies, combining experimental work with mathematical methods has been a successful approach to gain quantitative insights into Th cell dynamics and decision-making (9–15). Notably, it was found that although signal integration via cytokines is transient and stochastic (16,17), the resulting decisions regarding the generation of T cell phenotypes, including selective cytokine secretion, are remarkably stable even in quantitative terms at the single-cell level (10). Nevertheless, assessing the complex interplay of different regulatory elements shaping the phenotypic Th cell landscape has been exacerbated by the limited availability of kinetic data, which are difficult to obtain experimentally because of small cell numbers occurring in vivo especially at early time points. Indeed, experimental and theoretical studies have underlined the value of time-course information for the quantitative understanding of dynamic processes such as T cell differentiation (4,18–25).

A still unresolved question in Th cell differentiation is the lineage identity of mixed cell phenotypes such as Th1/2 hybrid cells. Those cells stably co-producing T-bet and GATA-3 have initially been discovered to arise in mouse models of parasite infections (7), their development was successfully recapitulated in vitro (7,17), and they are a common observation in recently available single-cell phenotyping data sets (8,26). Other non-conventional Th cells comprise Tfh-like PD-1^hi^CXCR5^-^, ‘peripheral helper’ T cells in rheumatoid arthritis (27), and Th17 cells in a ‘poised type 2 state’ in the context of tissue injury (28). How do hybrid Th cell lineages relate to the conventional Thx lineages? In particular, do hybrid cells result from mixed or superimposed gene expression programs of two or more conventional lineages, for instance as a combination of genes driven by T-bet and GATA-3 transcription factors in the case of Th1/2 hybrid cells? Or, do they rather evolve toward independent gene expression programs during differentiation?

To address such questions, and to derive a comprehensive picture of transcriptional dynamics during Th cell differentiation, we performed a high-resolution kinetic analysis of gene expression changes with a 3 hr time interval for the very first time points. We followed Th cell differentiation into Th1 and Th2 cells, complemented by Th0 conditions and a Th1/2 hybrid phenotype, each in two independent kinetic transcriptomics experiments. We developed a quantitative workflow to carefully characterize the temporal expression patterns of kinetic genes, and to analyze differences between cell types arising in the kinetic transcriptional program. We found a critical lineage bifurcation point at ∼24 hrs after antigen stimulation. Notably, we identified a set of genes that show independent behavior in the Th1/2 hybrid cells and are under direct control of STAT1/4 rather than following T-bet– or GATA-3– dependent transcriptional programs.

## Results

### High-resolution kinetic gene expression analysis reveals a critical bifurcation point early during differentiation

Previous experiments have shown that Th cells can exhibit distinct and mixed phenotypes based on the combination of polarizing cytokine signals. Here, we used an established in vitro protocol combining T cell receptor (TCR) stimulation and polarizing cytokines, to induce Th cell differentiation towards Th1, Th2 and Th1/2 hybrid cells, supplemented by a Th0 condition with TCR stimulation and blocking antibodies for IFN-γ, IL-12 and IL-4 (Figure 1A)(7). The obtained Th cell lineages were analyzed by flow cytometry, indicating lineage-specific expression profiles of key cytokines and transcription factors, as expected (Figure 1B, Figure S1). In particular, Th1 cells showed a dominant T-bet and IFN-γ expression profile, Th2 cells showed GATA-3 and IL-4 expression, and Th1/2 hybrid cells showed a mixed phenotype. Th cell transcriptomes were obtained at 10 time points over a time-course of 120 hours, the first three time points in 3 hrs intervals. Two independent experiments were performed, with very similar overall data quality and gene expression kinetics. For many genes that are known to have an important role in Th cell differentiation, we observed strong up- or down-regulation within the time window of the experiment in a cell-type specific manner (Figure 1C, Figure S1A). As expected, genes of the well-known Th1 and Th2 signature cytokines and transcription factors, *Tbx21, Gata3, Ifng, IL4*, showed a cell-type specific early response in the corresponding polarizing conditions (Figure 1D). Further, the hybrid Th1/2 phenotype featured elevated expression levels of both *Tbx21* and *Gata3*, while Th0 cells showed *Tbx21* dynamics similar to Th2 cells and *Gata3* dynamics similar to Th1 cells.

**Figure 1:**
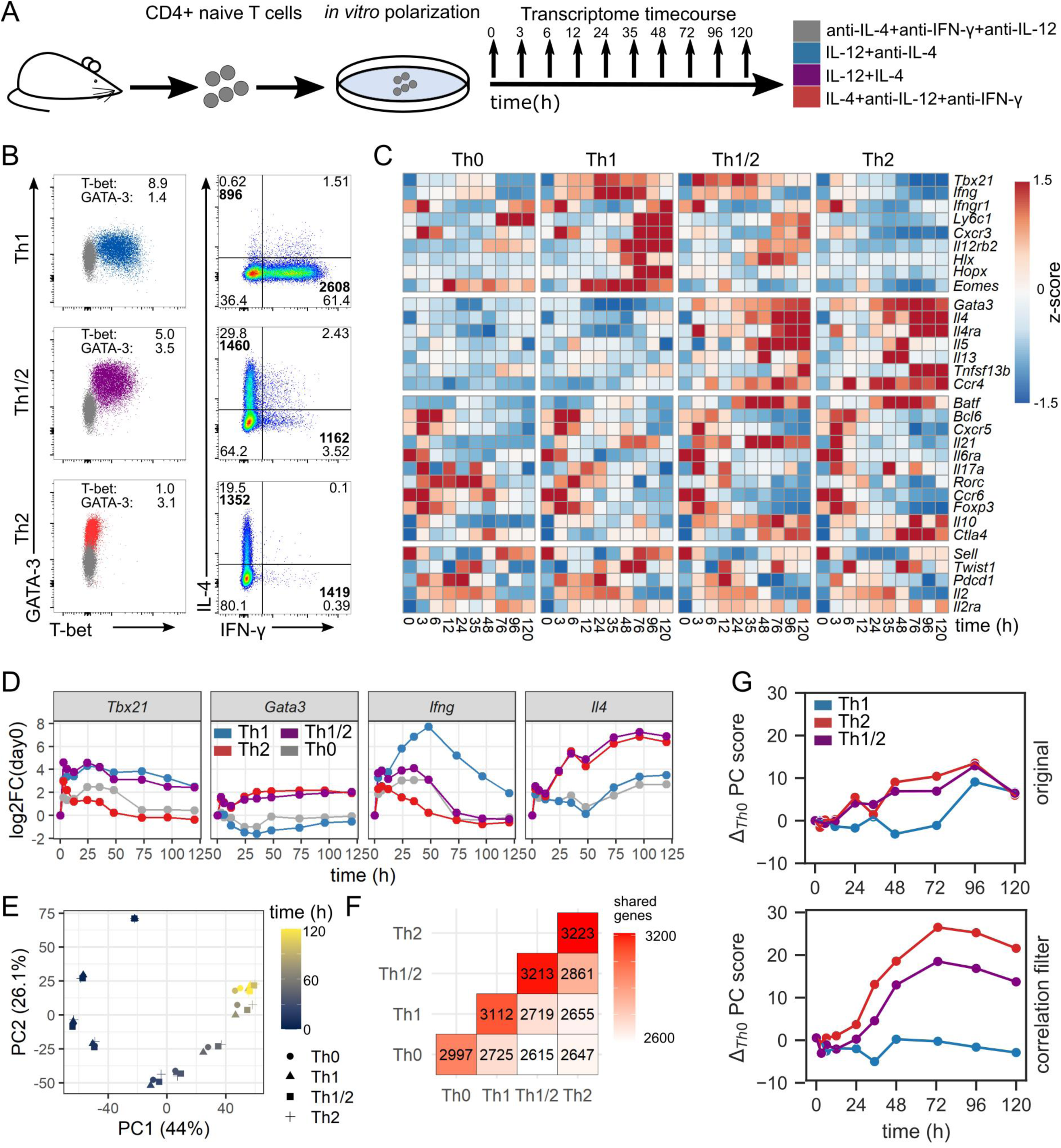
A high-resolution time course of Th cell differentiation. (A) Experimental setup. Th cell subsets were induced by polarizing signals in vitro, and gene expression profiles were obtained at 10 time points between 0 and 120 hours after activation. (B) Flow-cytometric characterization of Th cell subsets 120 hours after activation with polarizing conditions as described in (A). Normalized geometric mean indices for T-bet and GATA-3 expression are shown. Geometric mean intensities for IFN-γ and IL-4 positive cells are indicated in bold. (C) Gene-expression profiles of four groups of genes (top to bottom): Th1-related, Th2-related, Tfh and Th17-related, and other important Th cell-related genes. (D) Kinetics of master regulator transcription factors and signature cytokines for individual CD4+ T cell subsets. Shown are normalized expression intensities as fold-change relative to the first measured timepoint (0h). (E) Principal component (PC) analysis of the differentiation time course. Cell subsets are indicated by marker shape. Time of measurement is indicated by color. (F) Numbers and overlap of kinetic genes between cell subsets. (G) Evolution of PC1 over time. Genes with high correlation between subsets were removed (bottom) or kept for comparison (top).

To derive a first overview on general characteristics of the obtained data, we performed principal component analysis (PCA) and hierarchical clustering (Figure 1E, Figure S2B and C). Differences between the analyzed cell types increased gradually, and time was the variable accounting for most of the variance (Figure 1E). That is in line with our result of 3.944 kinetic genes out of 12.479 expressed genes obtained by a combination of statistical tests (cf. Methods) (Figure 1F). Next, to analyze the kinetics of cell differentiation, we removed genes that were highly correlated across all 4 subsets from the data set (Figure S2D). In a PCA on that reduced data set, differences between cell fates were far more pronounced than in the original data set (Figure 1G). The differences between cell fates started increasing after approximately 24 hours and reached a stable maximum at ∼day 3, which was consistent across all first four principle components (Figure S2E). Intriguingly, the Th1/2 hybrid cell type showed a deviating transient behavior in higher-order principle components (Figure S2E), already pointing to qualitative differences in the regulation of a fraction of genes that we shall explore in more detail below.

In summary, our explorative analysis of kinetic gene expression during Th cell differentiation revealed a bifurcation between individual cell types between day 1 and day 3, suggesting a critical time window for Th cell differentiation around day 1 after TCR stimulation.

### Early Th cell differentiation features three major patterns of kinetic gene expression

Having obtained an overview about the global transcriptomic changes during Th cell differentiation, we next analyzed the genes with significant changes over time in more detail. For this purpose, we first used the established MasigPro (29) software package to cluster the kinetic genes of each subset (Figure 2A, cf. Methods). We identified three dominating temporal patterns or kinetic clusters (Figure 2B-D, Figure S3A-D): fast and transient up-regulation (C1), delayed and stable up-regulation (C2), and stable down-regulation (C3). The three kinetic clusters occurred in comparable abundance across all cell types, cluster C1 occurring with slightly lower frequency compared to clusters C2 and C3 (Figure 2C). Many well-known Th1 and Th2 cell fate-inducing genes were identified as kinetic, and were associated with kinetic clusters in a cell-type specific manner (Figure 2D). In contrast, genes associated with other Th cell lineages such as *Rorc and Il17a* (Th17) or *Pdcd1* (Tfh) did not show a significant kinetic response according to our criteria. Finally, we performed pathway overrepresentation analysis for the kinetic genes associated to each cluster (Figure 2E, Table S1). We found that the stably up-regulated dynamics of cluster C2 were strongly associated with cell-cycle activity and metabolism, while the transient dynamics of cluster C1 showed enrichment for regulation of transcription and translation. Moreover, we identified early responses for type I interferons and IL-2 signaling in cluster C2, while other immune cell-related signaling activity was found throughout all clusters including the down-regulated genes in cluster C3.

**Figure 2:**
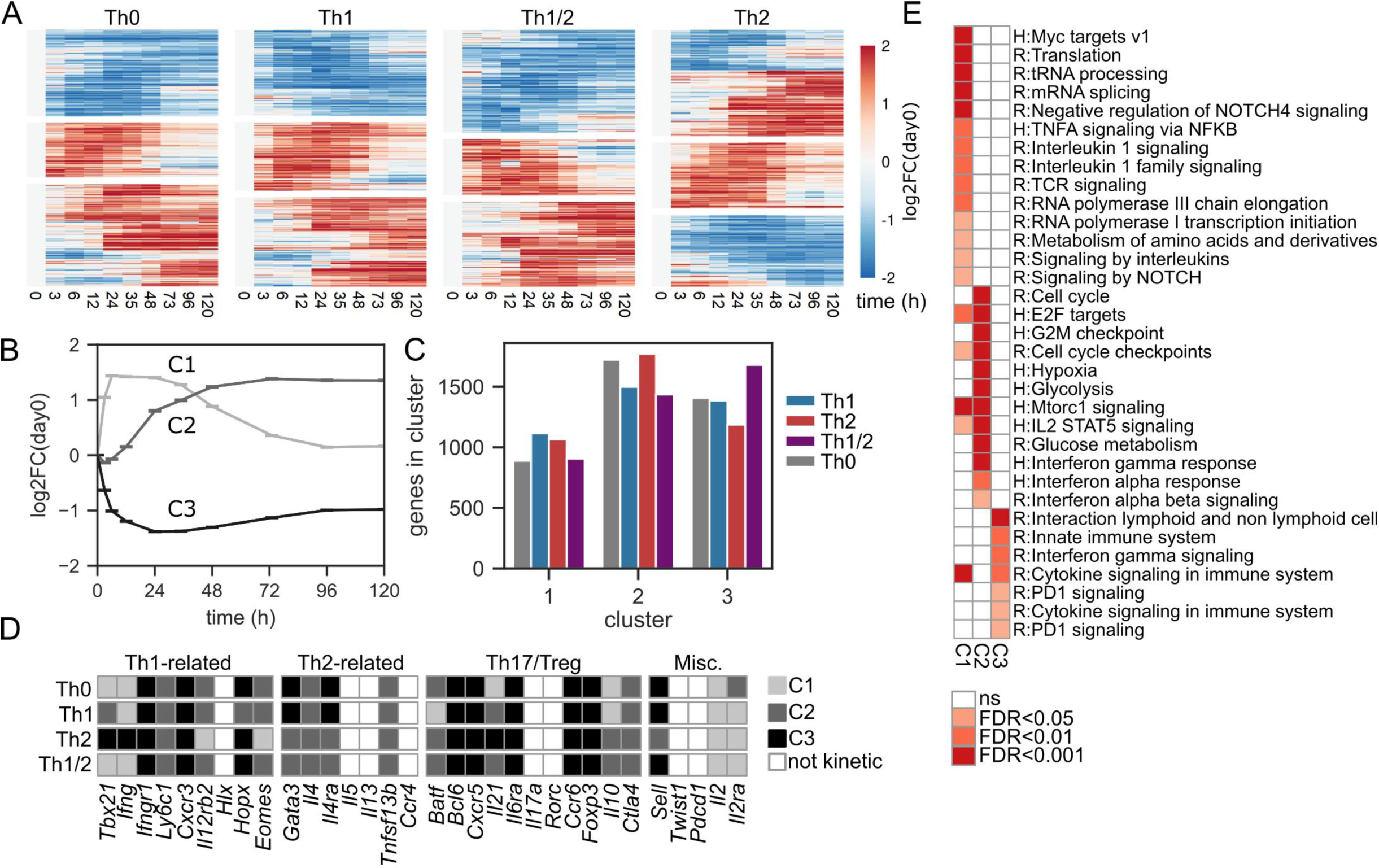
Early Th cell differentiation features three major patterns of kinetic gene expression. (A) Expression heatmap for kinetic genes. (B) Normalized expression kinetics of the three identified kinetic gene expression clusters, shown as averages over all genes and all cell types contained in each cluster. (C) Quantification of the numbers of identified kinetic genes across cell types within each kinetic cluster. (D) Gene classification as non-kinetic or kinetic incl. cluster association, for the four groups of Th cell-related genes introduced in Figure 1C. (E) Pathway enrichment analysis for genes uniquely assigned to kinetic clusters C1-C3. Pathways were pooled from REACTOME and Msigdb:Hallmark data bases, for a list of all enriched pathways see Table S1.

### A refined selection procedure identifies quantitative and qualitative differences in kinetic gene expression between Th cell subtypes

Based on the described set of kinetic genes, we next analyzed differences in the dynamics between cell types. To this end, we used a combination of the kinetic differentially expressed genes (DEG) as derived from the Masigpro workflow (quantitative DEG) and an additional filtering step to exclude genes with strong pairwise correlation over time (qualitative DEG) (Figure 3A). The latter approach allowed us to select for genes that not only show distinct expression levels over several time points, but also show dissimilar trends over time (Figure 3B). This approach is analogous to a “Volcano plot” representation, which is often employed for selection of genes with high fold-increase in static gene expression analysis workflows. Finally, we added a category “cluster switch” based on whether a gene was assigned to a different kinetic cluster (cf. Figure 2B) for each comparison of cell types.

**Figure 3:**
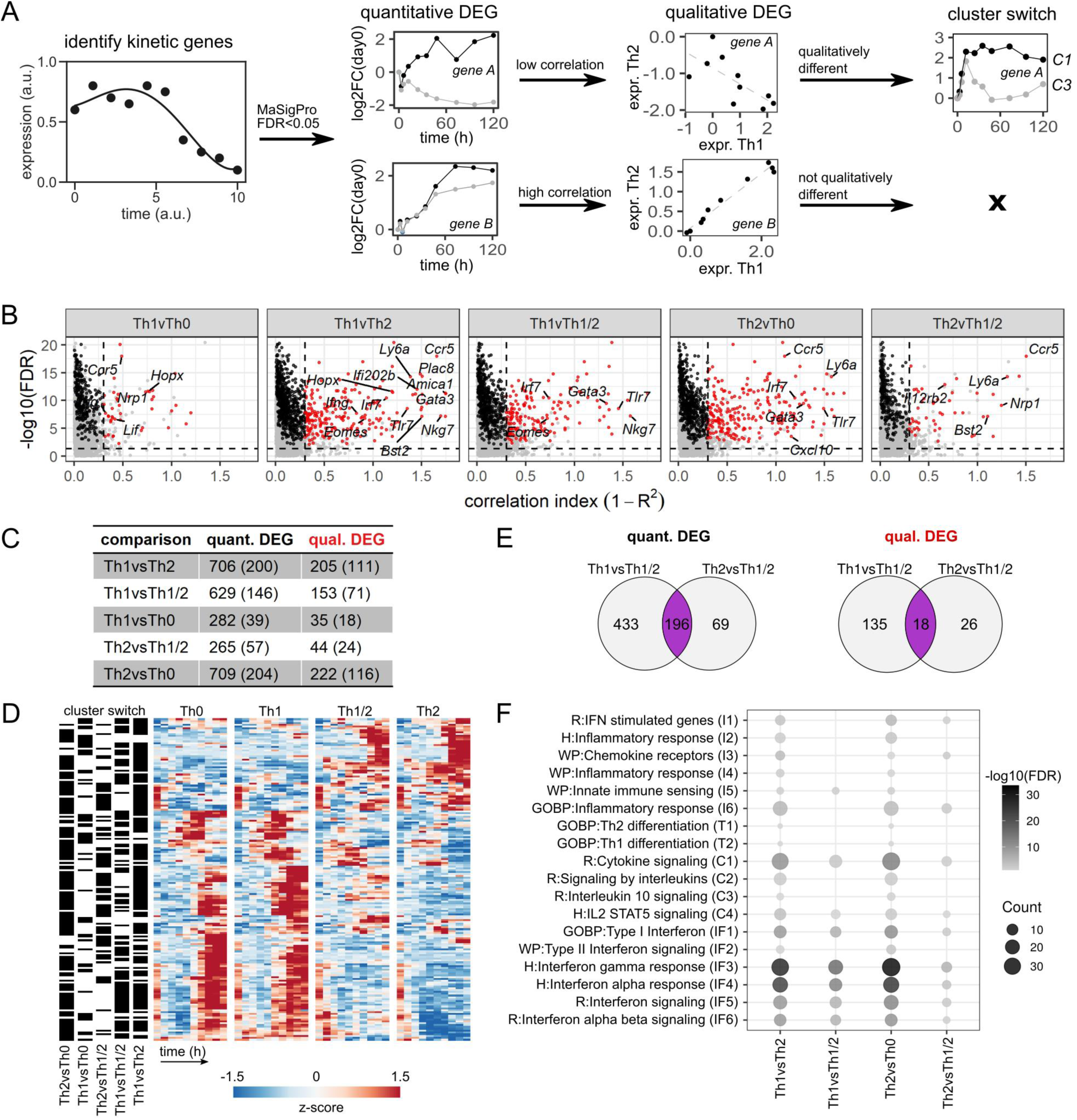
A refined selection procedure identifies quantitative and qualitative differences in kinetic gene expression. (A) Workflow illustration. We employed a combination of regression fitting in MasigPro to derive quantitative differentially expressed genes (DEG), followed by a correlation filter to identify qualitative DEG and by an analysis of switching of kinetic clusters between cell types. (B) Correlation volcano-plots based on the workflow in (A). Genes are categorized as kinetic (grey), quantitative DEG (black) or qualitative DEG (red). See Methods for details. (C) Numbers of qualitative and quantitative DEG obtained for each comparison of cell types. Brackets indicate the numbers of kinetic cluster switches. (D) Expression heatmap for all qualitative DEG exhibiting kinetic cluster switches in at least one comparison of cell types, as indicated on the left. (E) Venn diagrams of quantitative (left) and qualitative (right) DEG shared between Th1 or Th2 cells and Th1/2 hybrid cells. (F) Pathway enrichment analysis of DEG for all pairwise comparisons between cell types. Pathways were pooled from REACTOME, GO:BP, Msigdb:C2:WikiPathways and Msigdb:C3:TFT data bases. Shown are pathways with significant enrichment in at least two comparisons.

The set of kinetic DEG derived from our data set contained 706 quantitative DEG, out of which 205 are also qualitative DEG, out of which 111 also are subject to cluster switch, as exemplified for the Th1 vs. Th2 comparison (Figure 3C, Table S2). Visual inspection of this set of genes showed clearly distinguishable patterns between Th1 and Th2 cells (Figure 3D). Apart from Th1 vs. Th2 DEG, we found the highest numbers of DEG in the Th1 vs Th1/2 and Th2 vs Th0 comparisons (Figure 3C), as expected based on PCA analysis (cf. Figure 1E). Notably, we consistently identified DEG that were shared between the Th1 vs Th1/2 and Th2 vs Th1/2 comparisons, across quantitative, qualitative and cluster-switching DEG (Figure 3E), suggesting that not all parts of the Th1/2 cell transcriptome directly follow either the Th1 or Th2 cell gene expression program. As in the kinetic cluster analysis above, we found that many of the well-known Th1 and Th2 cell-associated genes such as *Gata3, Ifng, Eomes* and *Il4* were identified as DEG, supplemented by other genes such as *Nkg7* and *Bst2* (Figure 3B, Figure S3E, Table S2). Pathway overrepresentation analysis (Figure 3F, Table S1) revealed strong enrichment of interferon-related pathways (IF) across all comparisons, except for the Th1 vs. Th0 contrast, which did not contain any enrichment for the pathways we considered. T cell differentiation (T) and most of the pathways accounting for chemokine signaling and generic inflammatory patterns (I) were moderately enriched in the Th1 vs. Th2 and Th2 vs. Th0 comparisons only. The broader “cytokine” category (C) contained highly enriched pathways across all comparisons, but also pathways lacking significant hits for the Th1vsTh1/2 and Th2vsTh1/2 comparisons.

Overall, this high-resolution kinetic data set allowed for a fine-tuned approach to kinetic gene expression analysis in terms of quantitative, qualitative and kinetic cluster-switching DEG, yielding a quantifiable classification suitable for direct assessment of the role of each gene in lineage-specific Th cell differentiation programs.

### Hybrid Th1/2 cells are partly driven by a STAT1/4-dependent gene expression program that is independent of Th1 and Th2 cell specific gene regulation

Our analysis consistently revealed an overlap of Th1 vs. Th1/2 and Th2 vs. Th1/2 DEG (Figure 3D). That suggests that the majority of the kinetic transcripts in Th1/2 hybrid cells follows either the Th1 or the Th2 cell gene expression program, while a substantial fraction of the transcriptome differs from that of both Th1 and Th2 cells. We reasoned that such transcriptional kinetics could result from either “superposition”, that is a combined effect of Th1 and Th2 cell types of gene regulation, or from an “independent” gene expression program, that is an expression pattern that cannot be attributed to Th1 or Th2 cells nor to their combination.

To further investigate the relation of Th1/2 hybrid cells to Th1 and Th2 cells, we restricted the analysis to the set of Th1 vs. Th2 DEG, thereby focusing on genes that are highly related to differential Th cell fate-development (Figure 4A). Next, we set up a linear regression model to describe the transcriptional program of Th1/2 hybrid cells as a function of Th1 and Th2 cell gene expression. The resulting regression coefficients β_Th1_ and β_Th2_ for each gene span a plane in which additive and subtractive effects relating to Th1 and Th2 cell gene expression are directly accessible (Figure S4A). We grouped all considered Th1 vs. Th2 DEG into “Th1-like”, “Th2-like”, “Superposition” and “Independent” categories, based on the significance of the β_Th1_ and β_Th2_ regression fitting (Figure 4B and C)(cf. Methods). As expected based on the PCA and DEG analysis results, a large fraction of genes in the Th1/2 hybrid cell expression profile was classified as “Th2-like”, again indicating the overall similarity of the Th1/2 hybrid phenotype to the Th2 cell type (Figure 4C). Another large fraction of genes was classified as “Superposition” or “Independent”, and quite remarkably, we found those two categories at almost the same frequency.

**Figure 4:**
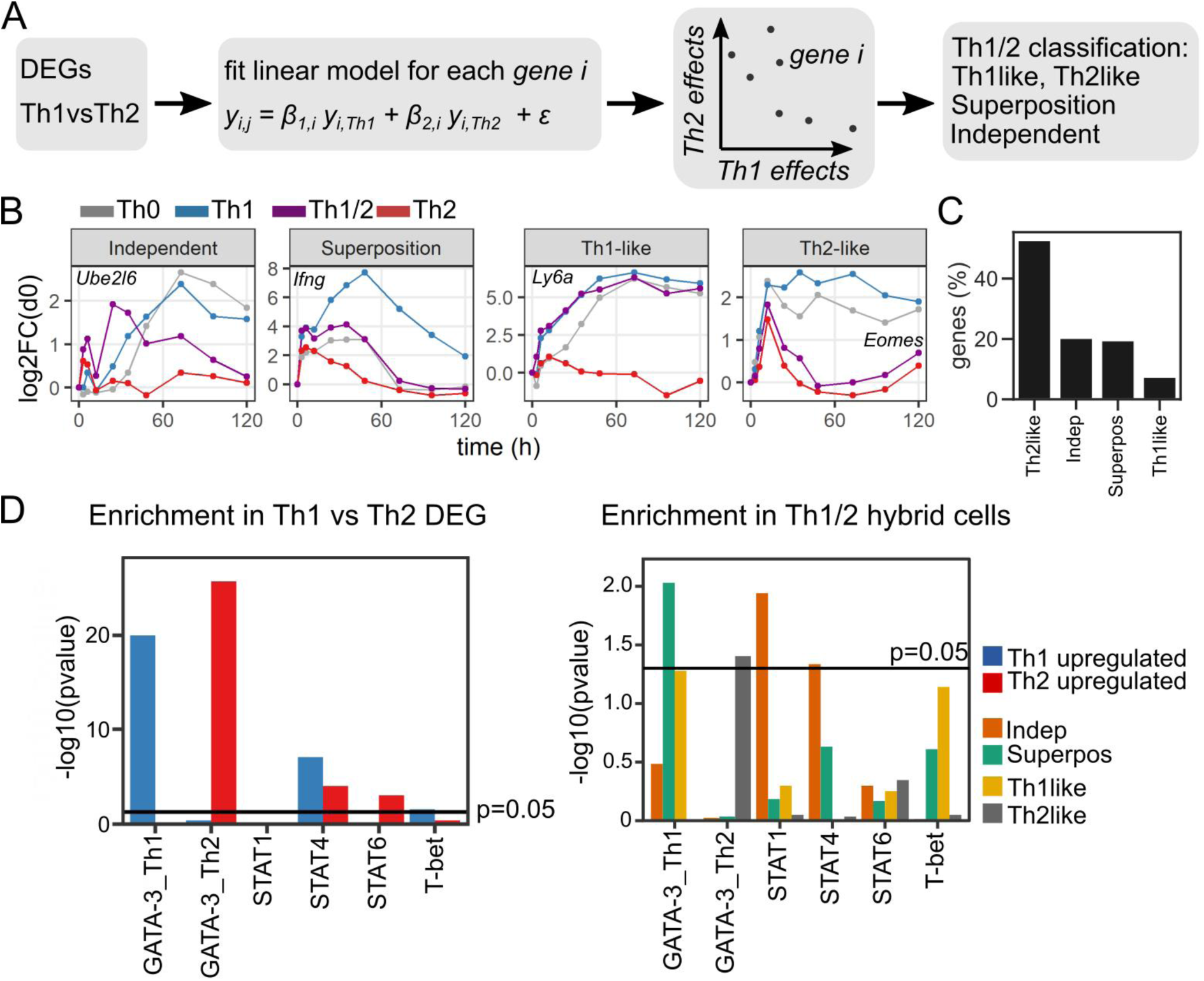
Superposition and independence of genes in Th1/2 hybrid cells. (A) Workflow sketch. Based on the identified qualitative Th1vsTh2 DEG (see Figure 3C), similarity of genes to the expression profile of Th1/2 hybrid cells was assessed by a linear regression model. (B) Time-courses of representative genes, and (C) quantification of gene classification into the four different categories. (D) Enrichment analysis using published transcription factor target gene sets (see text). Left: Analysis of up-regulated genes in Th1 (Th2) cells, which are taken as Th1vsTh2 DEG with expression values higher (lower) in Th1 compared to Th2 cells. Right: Analysis of the Th1/2 hybrid transcriptional profile along the gene categories obtained in (A-C).

To further evaluate the described types of genes in context of the overall transcriptional program, we performed enrichment analysis with regard to publicly available gene lists of transcription factor targets as obtained from ChipSeq data (30–34). We focused on gene regulation by the transcription factors GATA-3, T-bet, and STAT1/4/6, which are known to be key regulators of Th cell differentiation. As expected, in the overall Th1 vs. Th2 contrast, the Th1 cell-related genes were enriched for T-bet, STAT4 and Th1-cell specific GATA3 targets, while the Th2 cell-related genes were enriched for STAT6 and Th2-cell specific GATA-3 and targets (Figure 4D, left panel). In the Th1-like and Th2-like genes of the Th1/2 hybrid cells, we also found strong enrichment for T-bet and GATA-3 target genes, respectively (Figure 4D, right panel). The superposition genes showed strong enrichment in the GATA-3 target genes of Th1 cells. In contrast, in the Independent genes of the Th1/2 hybrid cells, we identified a significant signature of STAT1 and STAT4 target genes that is absent in all other types of Th1/2 hybrid cell genes. This pattern of a dominating STAT-dependent transcriptional program for Independent genes and dominating GATA-3-dependent regulation for Superposition genes was consistent for our two independent replicates and was robust to changes in the applied thresholds for statistical analysis (Figure S4B-E).

Taken together, we found that the majority of genes in the Th1/2 hybrid cells closely follow either the Th1 or Th2 cell transcriptional programs, but about 20% of the remaining genes showed independent behavior rather than being explained by a combination of Th1- and Th2-dependent effects. In contrast to the expression profiles of Th1 and Th2 cells, which were dominated by T-bet and GATA-3 control, those independent genes in Th1/2 hybrid cells were primarily comprised of STAT1 and STAT4 target genes.

## Discussion

The commitment of Th cells to a specific effector state is one of the key decision-making processes at the beginning of an immune reaction and has far-reaching consequences regarding the type and strength of the response. That decision can have severe consequences in the context of diseases including autoimmune disorders (35,36), cancer (37), or viral infections including SARS-CoV-2 (38). Here, kinetic gene expression analysis at high temporal resolution especially in the very early phase of cell differentiation allowed us to derive a full picture of the transcriptional landscape during Th1, Th2 and Th1/2 cell differentiation, and to achieve a detailed classification of the kinetically changing genes. We could pinpoint a critical time window at ∼24 hrs after TCR stimulation, where the lineages start to show divergent behavior, and we provide detailed information regarding kinetic patterning of genes within and between Th cell effector subtypes.

Recently, high-content single-cell technologies such as CyTOF and single-cell sequencing have allowed deep insights into the rich and previously unforeseen diversity of the phenotypic space of effector Th cells, which can cover the full spectrum between and around the conventional Th1, Th2, Th17 etc. cells (8,26,39,40). Furthermore, non-conventional Th cell phenotypes have been discovered for instance in the contexts of rheumatoid arthritis and tissue injury (27,28). Such findings have raised the question whether immune cell phenotypes should be regarded as a continuous landscape rather than a set of discrete states (40). Here, using kinetic transcriptome analysis after highly controlled generation of Th1, Th2 and hybrid Th1/2 cells in vitro, we were able to follow the gene expression dynamics in all three cell types simultaneously. In particular, we could directly compare the changes of individual genes between the hybrid cells and the related conventional Th1 and Th2 cells over the full time-course of Th cell differentiation.

We found that despite the co-expression of T-bet and GATA-3 in the Th1/2 hybrid cells, the majority of genes showed “bi-modal” behavior and closely followed either the Th1 or the Th2 cell type dynamics. Only a fraction of ∼20% of genes showed the expected “in-between” behavior, that is, a superposition of the Th1- and Th2-dependent effects. An equal portion of again ∼20% of genes showed independent behavior, that means the temporal evolution of those genes could not be attributed to either Th1 or Th2 kinetic patterns or the combination of both. Notably, we found that the independent genes in the Th1/2 hybrid cells do not follow the otherwise dominant signature of T-bet or GATA-3 target gene enrichment, but rather are under the control of STAT1- and STAT4-dependent gene regulation. Such dominant STAT1/4 control might be a consequence of GATA-3 and T-bet dependent gene regulation cancelling the effect of each other in those genes.

Hence, our analysis revealed substantial commitment of the hybrid Th1/2 cell lineage to the corresponding conventional, polarizing Th1 and Th2 cell lineages; nevertheless, we also identified fractions of the gene expression program accounting for independent or intermediate states. That suggests that the question of a continuous versus discrete gene expression landscape of Th cell lineages depends on the individual gene or gene set under consideration. Here, deep time-course transcriptomic profiling generated a resolution allowing for such detailed analysis of the phenotypic identity among closely related immune cell types.

## Materials and Methods

### Mice

Balb/c mice were bred under specific pathogen-free conditions at the Charite, Berlin. All animal experiments were performed in accordance with the German animal protection with permission from the local veterinary offices.

### Cell culture and in vitro differentiation

Cells were isolated and cultured as previously described (7). Briefly, naïve CD4+ CD62L^hi^ T cells were isolated from pooled spleen and lymph node cells of 5-8 week old Balb/c mice using a two-step magnetic sorting strategy (Multisort kit, Miltenyi Biotec). T cells were cultured in RPMI 1640+GlutaMax-I supplemented with 10% (v/v) FCS (Gibco), penicillin (100 U/ml; Gibco), streptomycin (100 µg/ml; Gibco), and ß-mercaptoethanol (50 ng/ml; Sigma). Cultures were prepared by stimulation with plate-bound 2.5 µg/ml anti-CD3ε (145-2C11) and 3 µg/ml soluble anti-CD28 (37.51, both from BD Biosciences). For Th1 differentiation, 10 ng/ml IL-12 (R&D Systems), and 10 µg/ml anti– IL-4 (11B11) were added. For Th2 differentiation, 30 ng/ml IL-4 (R&D Systems), 10 µg/ml anti–IL-12 (C17.8), and 10 µg/ml anti–IFN-γ (AN18.17.24) were added. Hybrid Th1/2 cells were cultured with 10 ng/ml IL-12, and 30 ng/ml IL-4. Th0 cells were generated under neutral conditions with anti–IL-12, anti–IFN-γ, and anti–IL-4. Cell cultures were transferred to a new plate and split on d 2. Transcription factor stainings were performed as previously described (7). T-bet and GATA-3 protein amounts were analyzed using FoxP3 staining buffer set (eBioscience) according to the manufacturer’s instructions. Briefly, cells were stained with anti-CD4 (RM4–5) followed by fixation with 1× Fixation/Permeabilization buffer and intracellular staining with PE-conjugated anti–T-bet (4B10) and Alexa-647–conjugated anti–GATA-3 (TWAJ, both from eBioscience) in 1× permeabilization buffer. Cells were washed in 1× permeabilization buffer and analyzed by FACS.

### Microarrays and data processing

Illumina microarrays (Illumina Mouse Sentrix-6) were used to profile T cell gene expression under polarizing conditions at 10 time points. Data were background corrected, quantile-normalized and log2-transformed. As an additional filtering step, we selected only probes whose expression was above the median expression across all groups and timepoints for at least one condition. Afterwards, we selected only probes that had gene annotations for EntrezGene ID, RefSeq ID and gene symbol, resulting in the analysis of 18284 probes (out of 46089), matching to 12479 expressed genes.

### Kinetic gene expression analysis

To identify kinetic genes, we first ran MaSigPro (29) on individual CD4+ T cell subsets to only consider time as an explanatory variable in the model. Additionally, we considered genes that had a 2-fold increase compared to time 0 at two consecutive time points. To identify temporal patterns, we employed hierarchical clustering using gene-gene correlation as a distance measure for each set of kinetic genes. The resulting dendrogram was cut at a prescribed number of clusters (Figure S4). Next, we used the union of all kinetic genes to also evaluate differences between different groups. To this end, we employed the MaSigPro workflow on all groups combined, thus considering time and group identity (Th1/Th2/Th12/Th0) as explanatory variables. To identify DEG over time, we first used a polynomial regression model as implemented in the MaSigPro package. Further, for the genes identified as having significant profile differences between groups (“quantitative DEG”), we computed for each gene the kinetic correlation using pairwise comparisons between cell types across all time points. Reasoning that a high correlation indicates a similar transcriptomic trajectory, we defined a correlation index 1-*R*^*2*^, where *R*^*2*^ is the Pearsson correlation coefficient, and identified all DEG with correlation index <0.3 between cell types as “qualitative DEG” (Figure 3A-C). For pathway analysis, gene sets were pooled from the public REACTOME, GO:BP, Msigdb:Hallmark and Msigdb:C23:Wikipathways data bases. We excluded gene sets with less than 3 or more than 1000 genes. Pathway overrepresentation analysis was performed by applying a hypergeometric test on gene sets of all databases combined after background correction.

### Linear model analysis

We used the following model to describe the expression for a gene *i* expressed in cell type *j* as a function of Th1 and Th2 expression: *Y*_*i,j*_ *= β*_*i,Th*1_*Y*_*i,Th*1_ + *β*_*i,Th*2_*Y*_*i,Th*2_ + *ε*. Here *Y*_*i,j*_ represents the gene expression for cell type *j ∈* {Th0,Th1/2}, and *Y*_*Th*1_ (*Y*_*Th*2_) the gene expression of Th1 and Th2, respectively. The coefficients *β*_*Th*1_ and *β*_*Th*2_ denote the contribution of the respective cells to explaining the expression for *Y*_*i,j*_. Fitting the model to each gene allowed classification into the categories Th1-like, Th2-like, Superposition and Independent, based on significance of the regression fit coefficients *β*_*Th*1_ and *β*_*Th*2_ (see Figure 4A and Figure S7A). Of note, some fits showed a mixed combination of positive and negative coefficients, which would indicate a combined effect of negative and positive regulation. However, in all those cases the negative coefficient was not significant.

### Statistics

The p-values derived from MasigPro or other methods were corrected for multiple-testing using the Benjamini-Hochberg method, if applicable. The resulting false-discovery-rate (FDR) or simple p-value was regarded significant at a significance level of 0.05, except for pathway enrichment analysis, where we accepted values of FDR<0.1.

## Data availability

The R-scripts developed for kinetic gene expression analysis will be released alongside the GEO data repository upon publication of the manuscript.

