## Supplementary Figures for "High-resolution kinetic gene expression analysis of T helper cell differentiation reveals a STAT-dependent, unique transcriptional program in Th1/2 hybrid cells"

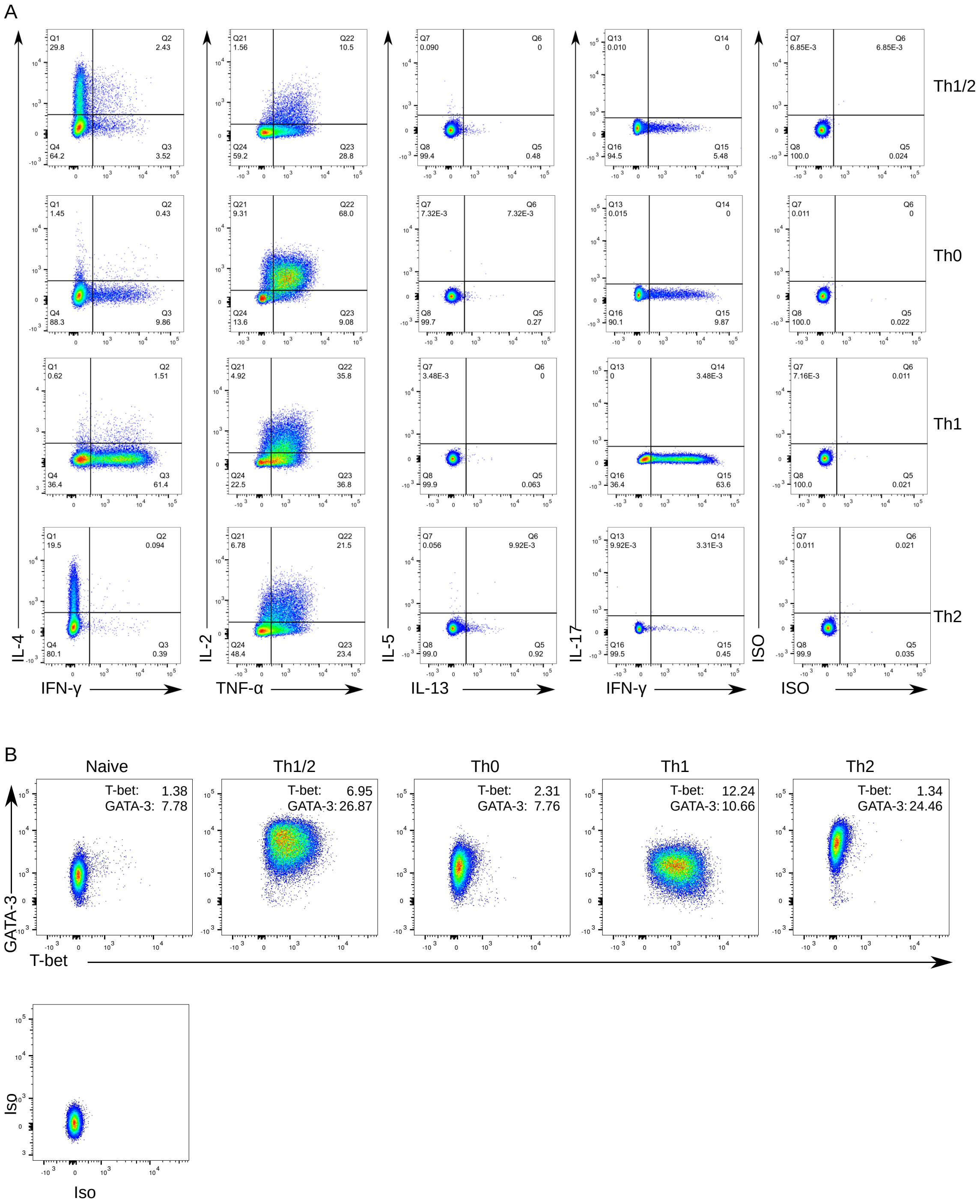


Figure S1: Flow-cytometric evaluation of Th cell subsets. (A) Flow cytometry analysis of signature cytokines of mouse-derived naïve CD4+ Th cells exposed to polarizing Th1, Th0, Th2 and Th1/2 conditions 120 hours after activation. (B) Staining for signature transcription factors of Th cell subsets for the same conditions as in (A). Geometric mean indices for T-bet and GATA-3 are shown.

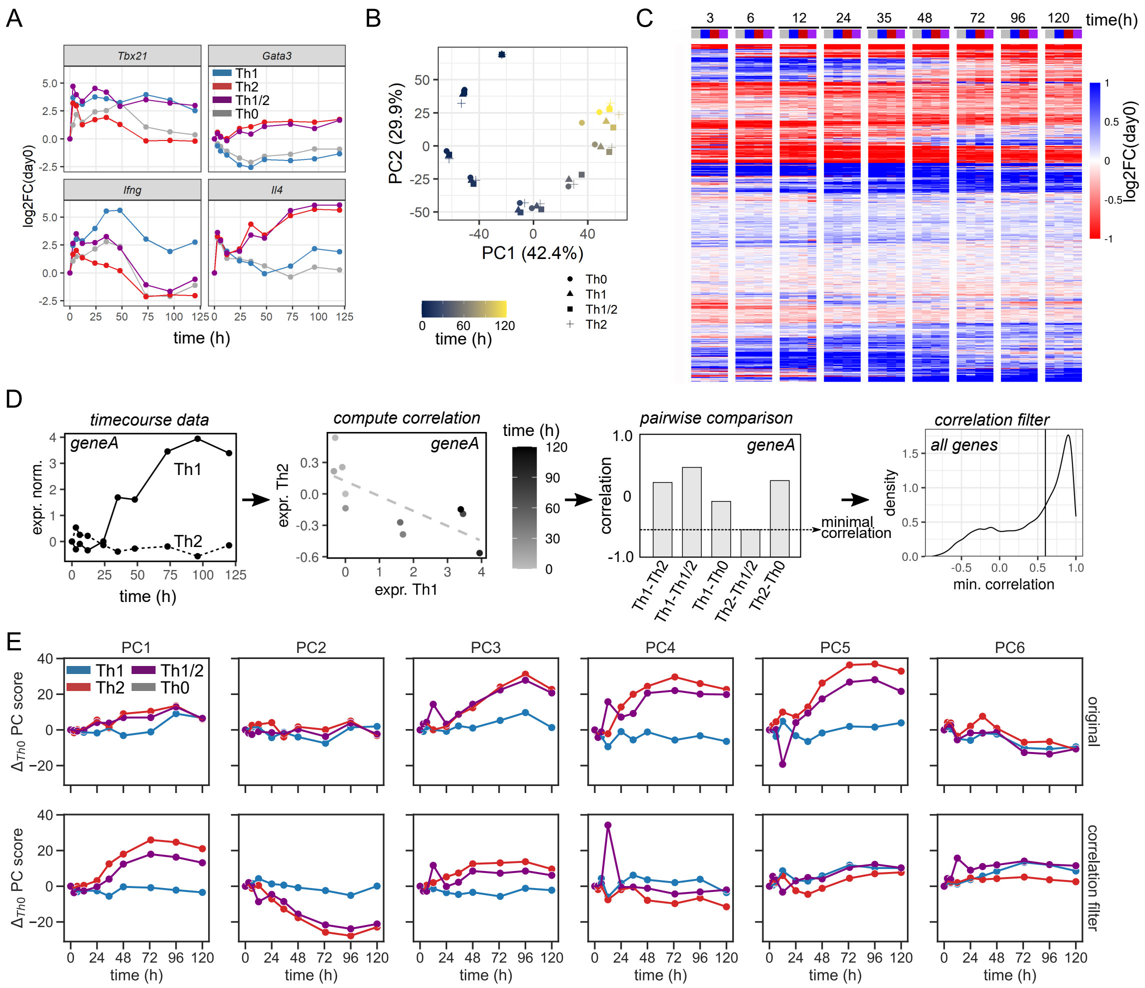

Figure S2: Exploratory data analysis and quality controls. (A) Transcription factors and signature cytokines for Th cell subsets in replicate 2. (B) PCA for replicate 2. (C) Heatmap of all expressed genes (replicate 1). Cell types are indicated by color as in panel (A). (D) Workflow to remove highly correlated genes. Gene-wise correlation was computed for all pairwise comparisons between samples. All genes with correlation coefficient exceeding a threshold value in at least one comparison were removed (cf. Methods). (E) Time-evolution of principal components with and without removal of highly correlated genes.

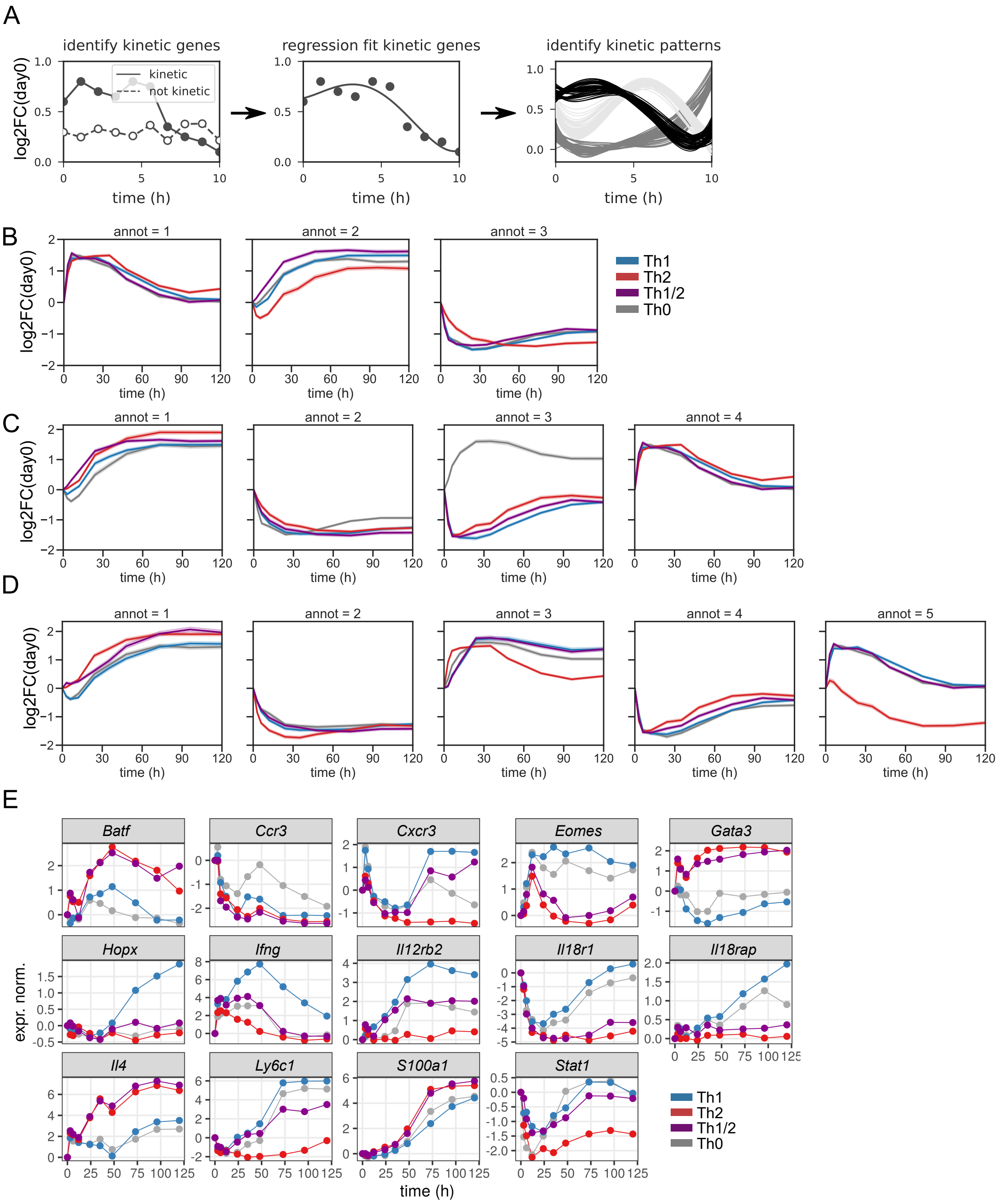


Figure S3: Supplementary analysis of kinetic gene expression profiles. (A) Regression-based MaSigPro workflow to analyze kinetic gene expression profiles . Kinetic genes are identified by regression and then clustered based on gene-gene correlation (cf. Methods). (B-D) Kinetic gene expression patterns computed setting the number of clusters to n=3,4 and 5. Shown are average gene expression values in each kinetic cluster for all cell types as indicated by color. (E) Expression kinetics of genes that exhibit cluster switches. Shown are genes with different kinetic cluster assignments between at least two cell types, and which are differentially expressed in at least one comparison (cf. Table S1).

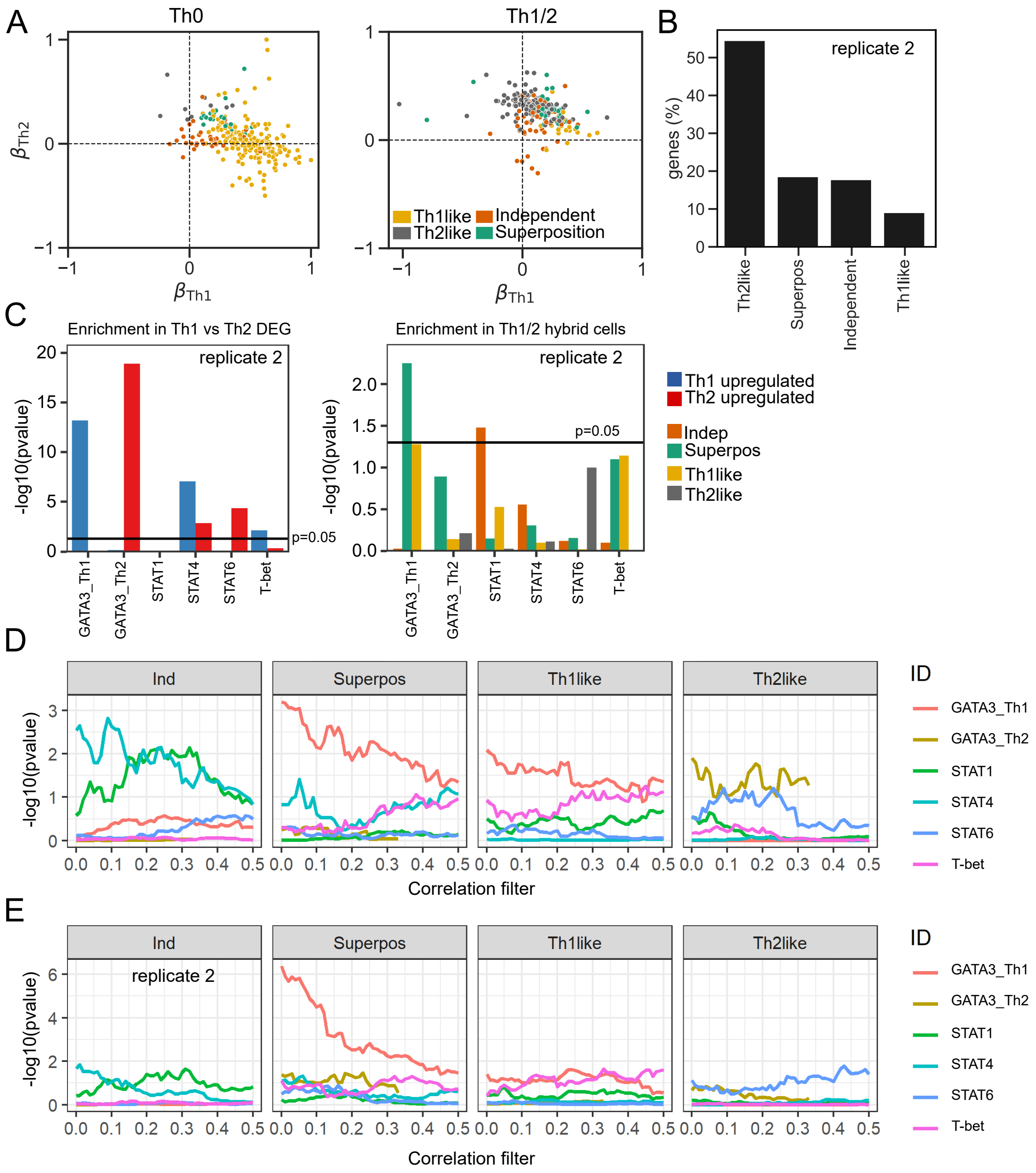


Figure S4: Supplementary analysis of gene expression profiles in Th1/2 hybrid cells. (A) Regression coefficients $\beta_1$ and $\beta_2$ of the linear regression model employed to identify superimposed, Th1like, Th2like and independent genes in Th0 and Th1/2 cells (cf. Methods and Figure 4A). Coefficients were normalized to the maximum value. (B) Category assignment into independent, superimposed, Th1like and Th2like genes for replicate 2 (cf. Figure 4C). (C) Enrichment results of transcription-factor target gene-sets for replicate 2 (cf. Figure 4D). (D-E) Enrichment of transcription-factor target gene-sets for different values of the correlation filter in replicate 1 (D) and replicate 2 (E) (cf. Methods).
